# Specialized cells and transporters mediate integument-absorption of environmental peptides in cichlids larvae

**DOI:** 10.1101/2024.10.19.619221

**Authors:** Pazit Con, Avner Cnaani

## Abstract

Nutrient absorption through the skin and gills into the organisms’ tissues has been documented in several aquatic invertebrates from different phyla. However, the actual absorption mechanism is still unknown. In teleost fish, as in all jawed vertebrates, intestinal absorption is considered as the sole source of nutrients. The proton-dependent peptide transporters (PepT) of the *slc15a* gene family are the only known mechanism for cellular absorption of di- and tri-peptides within the animal kingdom. In this study, we explored the expression and localization of PepT2 in Mozambique tilapia (*Oreochromis mossambicus*) larvae. Transcript levels of PepT2 in dissected yolk-sacs from larvae showed significant expression during the larval developmental period. Immunofluorescence staining of PepT2 with Na/K-ATPase (NKA) and Na^+^/K^+^/2Cl^-^ co-transporter (NKCC) on the yolk-sac membrane revealed co-staining with NKA and differential-staining with NKCC. While NKA staining was observed on the ionocytes’ basolateral membrane, PepT2 staining was restricted to the apical pit of the ionocytes, facing the surrounding water. In this study, we identified a nutrient transporter located on integument-specific cells, facing the outside aquatic environment. This is the first indication of environmental nutrients absorption in teleosts, and the first evidence of a possible absorption mechanism through PepT2, in specialized yolk-sac ionocytes.

## Introduction

Dissolved organic matter (DOM) can be a nutritional source, but usually support small organisms, such as bacteria or small invertebrates (Gomme, 2001). These organisms form the primary trophic levels of the food chain, initiating the flow of energy to higher-level consumers. Subsequent trophic levels, comprising larger and more complex organisms, depend on active predation to fulfill their nutritional requirements. These higher feeders preform active digestion of the feed into biochemical molecules, which are then absorbed along the digestive system. While this paradigm of feeding strategy is the most common, there have been several exceptions for higher trophic level aquatic invertebrates that can utilize DOM directly from the water. This phenomenon was described in an arthropod crustacean, with amino acids being absorbed through the posterior gills of the green shore crab (*Carcinus maenas*)(Blewett and Goss, 2017). The only known fish able to absorb nutrients through external integuments is the Pacific hagfish (*Eptatretus stoutii*), an ancient jawless vertebrate, in which, nutrients absorption was detected across both gill and skin surfaces (Glover et al., 2011). While these studies showed the absorption of environmental nutrients from the water into the organisms’ tissues, the actual absorption mechanism is unknown. Here, we demonstrate a novel localization of the transmembrane peptide transporter 2 (PepT2) at the integument of cichlids’ yolk-sac in the specialized ionocytes cells.

Within the gastrointestinal tract of multicellular organisms, ingested proteins are digested by enzymes to the final products: free amino acids or as short peptides. These smallest forms of proteins are taken up from the intestinal lumen by the enterocytes, one of the intestine epithelial cell types, through specialized membrane transporters, expressed at their apical membranes. These free amino acids and small peptides transporters are relatively conserved across the animal kingdom (Verri et al., 2016). Small peptides (i.e. Di- and tri-peptides) are recognized and transported by the proton-dependent peptide transporters (PepT), coded by the SLC15 genes family, which is one of the few known mechanisms for their cellular absorption. Higher Vertebrates has only two variants of this transporter, while in fish there are three variants: the PepT1a and PepT1b paralogs which are high-capacity/low-affinity transporters expressed in the intestine, and PepT2, which is a low-capacity/high-affinity transporter expressed in the intestine, kidney and brain of adult fish (Con et al., 2021). All three transporters are also express in the pre-feeding tilapia larva (Con et al., 2019).

Fish regulate ion levels in their body fluids through multiple tissues that allow interchanging ions with the surrounding water. As the integument acts as a barrier that limits water, gasses, and solutes exchange with the environment, specialized cells called ionocytes are responsible for ions removal or uptake and metabolic waste excretion (Evans et al., 2005). While in adult fish the ionocytes are located mainly in the gills, in larval stages the developing gills are not fully functional, and ion regulation activity is done mostly by ionocytes clustered on the yolk-sac epithelium (Kaneko et al., 2002; Katoh et al., 2000). In tilapia (*Oreochromis* spp.), four types of ionocytes were characterized on the yolk-sac, differing in their ion absorption or secretion capabilities (Hiroi et al., 2005).

## Results and Discussion

We used the quantitative real-time PCR and immunofluorescence staining (Con et al., 2019) to localize PepT2 on the yolk-sac membrane of Mozambique tilapia (*Oreochromis mossambicus*) larvae, and identified them in specific type of larval ionocytes. Transcripts levels of *slc15a2*, the gene encoding to PepT2, in dissected yolk-sac from larvae at 6, 9 and 12 dpf showed significant changes along the larval developmental period, with peak of expression at 9 dpf, following with decreased expression at 12 dpf (Figure 1A). Immunofluorescence staining of PepT2 and Na^+^/K^+^-ATPase (NKA) on whole larvae revealed co-staining of ionocytes on the yolk-sac membrane. While NKA was localized on the basolateral membrane of ionocytes, PepT2 staining was restricted to the apical pit of only few ionocytes. This was seen in larvae of both, Mozambique tilapia (Figure 1B) and Red belly tilapia (*Coptodon zillii*) (Figure 1C). The staining with Na^+^/K^+^/2Cl^-^ co-transporter (NKCC) antibody, showed basolateral staining of ionocytes sub-population, however, did not co-localized on the same cells that showed PepT2 apical staining (Figure 1F). These staining patterns seen with NKA and NKCC indicate that PepT2 is localized in the previously described Type-I ionocytes, a sub-population with unknown function (Hiroi et al., 2005), pointing to a potential larval-specific significant function of this transporter.

**Figure 1.**
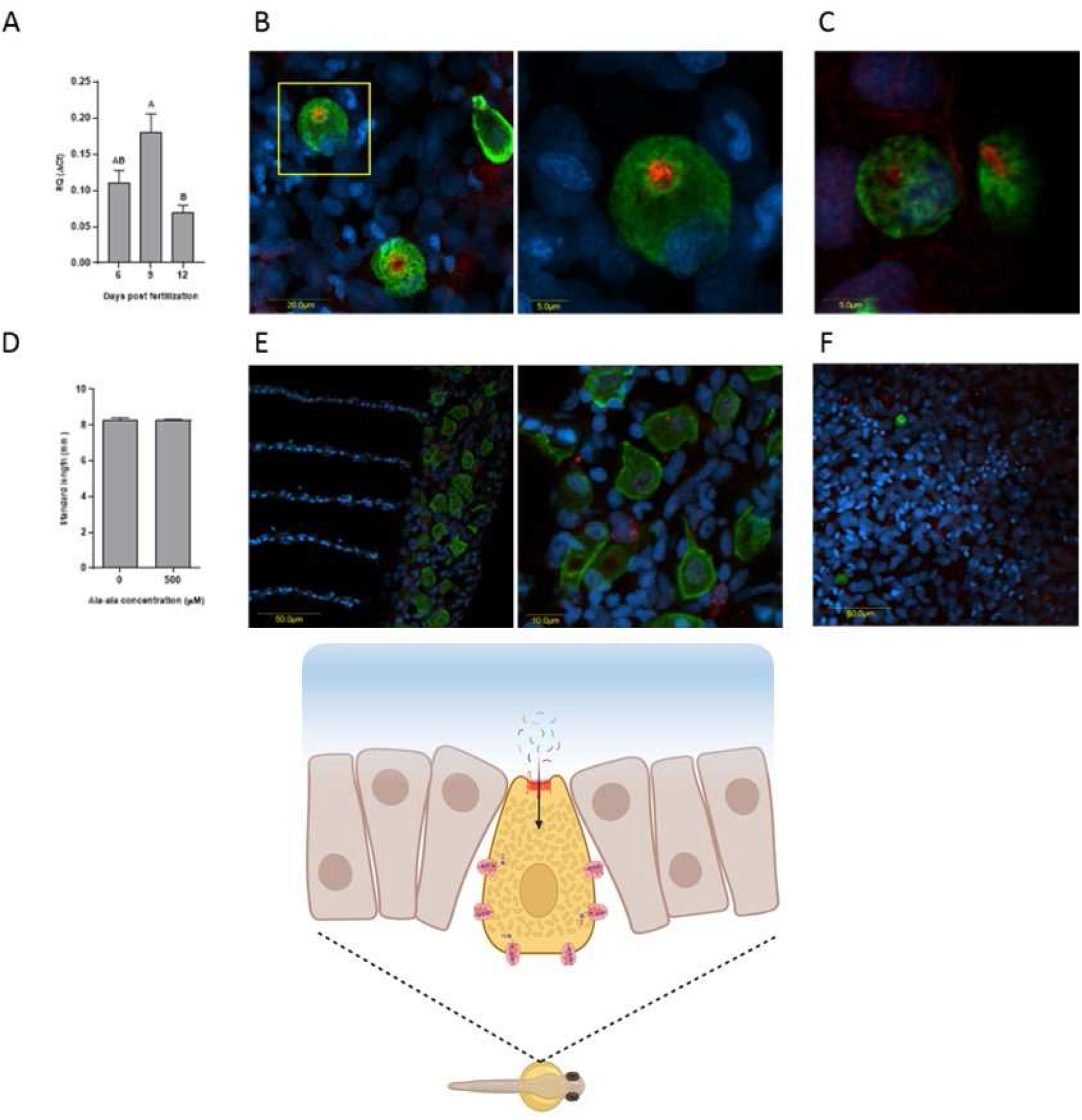
(A) qRT-PCR analysis of PepT2 in the yolk sac of *O. mossambicus* larvae at 6, 9 and 12 days post fertilization (dpf). (B) Immunofluorescence staining of PepT2 (Red) and NKA (green) with nuclei staining (blue) of 6 dpf *O. mossambicus* larvae yolk-sac ionocytes. (C) Immunofluorescence staining of PepT2 (Red) and NKA (green) with nuclei staining (blue) of 3 dpf *C. zillii* larvae yolk-sac ionocytes. (D) Standard length measurements of 14 dpf *O. mossambicus* larvae reared at different di-peptide concentrations. (E) Immunofluorescence staining of PepT2 (Red) and NKA (green) with nuclei staining (blue) of adult *O. mossambicus* gills. (F) Immunofluorescence staining of PepT2 (Red) and NKCC (green) with nuclei staining (blue) of 6 dpf Mozambique tilapia (*O. mossambicus*) larvae yolk-sac ionocytes.

These results demonstrate that in the tilapia larvae, PepT2 faces the surrounding water at the apical membrane of specialized ionocytes, located on the yolk-sac integument. While several aquatic invertebrates were shown to absorb nutrients across external epithelia, the actual cellular mechanisms of environmental nutrient absorption remained obscure. Here, we show the first indication of such absorption in jawed vertebrate, and the first evidence of the potential absorption mechanism.

The physiological relevance of environmental peptide absorbance in cichlid larvae is still unclear. In hagfish, that can survive several months between meals it was suggested that this phenomenon is important to maximize nutrient acquisition when immersed in decaying carcasses (Glover and Weinrauch, 2019). In tilapia larvae, however, the potential addition of nutrients from the surroundings is likely negligible compared to the nutritional resources of the large yolk sac which support the larval development for 10-14 days. This is supported by the growth trial results, which showed lack of growth differences between larvae reared in different concentrations of environmental peptides (Figure 1D).

Staining of gills sections from adult fish with PepT2 and NKA antibodies only showed positive staining of NKA at the basolateral membrane of ionocytes, but no PepT2 staining was found in gills ionocytes (Figure 1E), pointing to a potential larval-specific significant function of this transporter. Considering the parental care that characterize tilapiine fishes of the family Cichlidae, whether mouth-brooding like the *O. mossambicus*, or eggs grooming in substrate-spawners like the *C. zillii*, the relevance of the environmental peptide absorption to larvae of these fish species may suggest that this mechanism was developed as route to transfer molecules from parent to offspring. Other than mediating the cellular uptake of di- and tri-peptides, PepTs are also able to recognize and transport pharmacologically important compounds, including some amino-cephalosporins (β-lactam antibiotics), angiotensin-converting enzyme inhibitors and antiviral prodrugs (Smith et al., 2013). Additionally, it was described as a mediator for inflammation response through the absorption of bacterial-derived small peptides (Charrier and Merlin, 2006). The unique expression of PepT2 towards the water in cichlids larvae could be related to immune mechanism during the larval stages.

This study showed that the cichlids larvae express PepT2 in the apical pit of yolk-sac Type-I ionocytes, facing the surrounding water. It is the first to identify a nutrient-transporter facing the water at the epidermis, locate it at specific cells and discern a cellular pathway of peptides acquisition from the surrounding aquatic environment in a teleost fish.

## Materials and Methods

The breeding set-up and the collection of Mozambique tilapia larvae were as previously described by (Con et al., 2019). Larvae were sampled at days 6, 9 and 12 post fertilization (dpf), in order to track gene expression in the yolk sac. At sampling, larvae were moved to ice cold water and yolk sac membranes from 60 larvae were removed by microsurgery and pooled for six replicates (10 larvae in each pool). RNA extraction, cDNA synthesis, and primers used and quantitative real-time PCR analyses were done as previously described (Con et al., 2019).

For environmental amino acids concentration effects on growth, 120 larvae from a single spawn were split to 12 conical (6 for each treatment) glass bottles with 100 ml of artificial FW (0.05% Red-sea salt in DDW) with or without additional 500μM Ala-Ala peptide. The artificial FW of both treatment were filtered with 0.2μm nylon filter in order to eliminate bacteria form the water. The larvae were grown at 25°C on an orbital shaker at 70rpm for 14 days and water were replaced every two days. At 14 dpf, three larvae were sampled from each bottle, fixated with 4%PFA, washed with PBS and examined under Leica fluorescence binocular and standard length was measured.

Tilapia-specific rabbit αPepT2 and mouse αNKA antibodies were used for gills sections and whole-mount immunofluorescence staining modified protocol. Larvae fixation was achieved by 10 min PFA, followed with 16 hours incubation in 100% methanol at 4°C. Staining procedure was conducted as previously described (Con et al., 2017) for gills sections, and was modified for whole mount immunofluorescence adjustments. No antigen-retrieval was conducted in order to keep whole larvae structure. Three hours incubation was used for blocking stage (3 hours at 5% NGS, 1% BSA in PBS) and 48 hours for primary antibodies incubation at 4°C. The larvae and adult gills sections were examined using a confocal microscope.

This study was approved by the Agricultural Research Organization Committee for Ethics in Experimental Animal Use, and was carried out in compliance with the current laws governing biological research in Israel (Approval number: IL-650/15).

## ACKNOWLEDGMENTS

We are thankful to Eduard Belausov for his imaging expertise and to Tatiana Slosman for the care and maintenance of the fish. This research was supported by grant IS-4800-15 from the US-Israel Binational Agricultural Research and Development Fund (BARD) and by grant 2016611 from the US-Israel Binational Science Foundation (BSF).

## Author contributions

P.C. and A.C. designed the research; P.C. performed the research and analyzed the data; P.C. and A.C. wrote the paper.

